# Species-level classification provides new insights into the biogeographical patterns of microbial communities in shallow saline lakes

**DOI:** 10.1101/2023.12.06.570325

**Authors:** Polina Len, Ayagoz Meirkhanova, Galina Nugumanova, Alessandro Cestaro, Erik Jeppesen, Ivan A Vorobjev, Claudio Donati, Natasha S Barteneva

## Abstract

Saline lakes are rapidly drying out across the globe, particularly in Central Asia, due to climate change and anthropogenic activities. We present the results of a long-read next generation sequencing analysis of the 16S rRNA-based taxonomic structure of bacteriomes of the Tengiz-Korgalzhyn lakes system. We found that the shallow endorheic, mostly saline lakes of the system show unusually low bacterioplankton dispersal rates at species-level taxonomic resolution. The major environmental factor structuring the lake’s microbial communities was salinity. The dominant bacterial phyla of the lakes with high salinity included a significant proportion of marine and halophilic species. In sum, these results, which can be applied to other lake systems of the semi-arid regions, improve our understanding of the factors influencing lake microbiomes undergoing salinization in response to climate change and other anthropogenic factors. Our results show that finer taxonomic classification can provide new insights and improve our understanding of the environmental factors influencing the microbiomes of lakes undergoing salinization in response to climate change and other anthropogenic factors.

## Introduction

Lake ecosystems are among the most rapidly and extensively altered ecosystems and have shown major changes in physico-chemical topology and biotic characteristics in the recent past^1–3^. Sometimes referred to as “meta-systems”, biodiversity of lakes is strongly affected by lake connectivity, ecosystem structure and dynamics, and their relative position in the landscape^4^. The instrumental value of lakes as an indicator of Earth’s response to climate change^5^ makes lake research an essential component of the IPCC and UNFCCC agenda. Major consequences of climate change for lake ecosystems are observed worldwide and are likely to be amplified in the future due to, for example, changes in ice phenology, lake surface water temperature and evaporation^6^. This has significant implications for water level and water quality, nutrient dynamics and trophic structure^7^, community composition^8,9^ and susceptibility to invasive species^10^.

The globally projected change in temperature and precipitation patterns^11,12^ affects, in particular, regions with a semi-arid climate and constitute a major threat to the biodiversity and functionality of lake ecosystems here. Central Asia, a semi-arid region harboring the largest number of endorheic lakes^13^, is also one of the most rapidly warming regions of the world^14^. Increasing temperature and, as a result, precipitation/evapotranspiration imbalance can lead to salinization and desiccation of saline and freshwater terminal lakes^15,16^ and this may have major effects on ecosystem structure and functioning^17–19^. Among environmental gradients, salinity is known as a major factor driving the diversity and composition of microbial communities on a global scale^20^ and, specifically, in lake ecosystems^21^. However, our understanding of the impact of salinity and salinization processes is limited due to geographical and taxonomic bias in the current literature^22^. The authors highlight the lack of available data concerning small water bodies (i.e., shallow lakes and ponds), datasets from semi-arid and arid regions, and studies focusing on microorganisms rather than aquatic invertebrates.

Until recently, the field of microbial community analysis has been dominated by Illumina platforms that rely on partial 16S rRNA gene sequences (≤300 bp) for OTU generation and taxonomic classification. However, with the emergence of new high-throughput sequencing techniques, such as Nanopore and PacBio, which can produce full-length 16S sequences, it has been demonstrated that Illumina reads cannot achieve sufficient taxonomic resolution to accurately differentiate between bacterial taxa^23,24^. For analysis of microbiomes, the longer reads provide significantly improved taxonomic resolution to species or even strain-level^25,26^. The third generation sequencing technologies, such as nanopore-based sequencing by Oxford Nanopore Technologies (ONT), not only overcome these limitations, but also allow for sample multiplexing and metagenomic sequencing^24,27,28^. The only concern about nanopore-produced long reads - during its initial development stages - was the relatively high error rate^29^. However, besides continuously improving chemistry kits and basecalling algorithms, bioinformatic approaches are being developed to handle noisy data^30–33^.

Here, we implement an improved nanopore-based workflow to comprehensively characterize lake microbiomes at high taxonomic resolution. We investigated the diversity, heterogeneity, and detailed composition of prokaryotic communities of the Tengiz-Korgalzhyn Lakes system, in Kazakhstan, located along the north border of the endorheic basin of Central Asia. We hypothesize that environmental gradients (mainly salinity) and lake connectivity are key drivers of the variation in biodiversity and composition of microbial populations in saline lakes. In addition, we anticipate that the species-level taxonomic profiling of the bacterial full-length 16S amplicons would help us to gain new insights into the microbial ecology of the ecosystems of these endorheic lakes, specifically the importance of environmental selection and dispersal processes in shaping bacterioplankton communities of neighboring and distant lakes.

## Materials and Methods

### Study area and sampling site classification

The Tengiz-Korgalzhyn Lakes system (TKL) is located in the Korgalzhyn district, Akmola region, Northern Kazakhstan. The TKL area was included in the Ramsar convention in 1976 and later added to the “Living Lakes” list by the Global Nature Fund in the early 2000s. The territory is also partially designated as the Korgalzhyn State Nature Reserve, which is currently listed as one of the UNESCO World Heritage Sites. Despite the protection measures, TKL remains under the pressure of anthropogenic and environmental factors, such as the utilization of water resources by the nearby towns, fluctuating water levels due to the operation of connected water dams, seasonal floods, droughts, etc. The region is defined by its continental and arid climate, with relatively scarce precipitation during the summer^34^. Most of the lakes are snow-fed, with little to no reliance on local temporary water streams^35^. Coastal sampling (1-2 m from the coast, 0.5 m depth) was conducted across the TKL and in several adjacent water bodies (**Figure 1**). For geographical and environmental comparison of the samples, we defined several scales to appropriately address the samples: region (the lowest scale), lake, and site (the finest; each sample corresponds to a single site). Hence, the studied area was divided into five regions: Nature Reserve, North Group, South Group, East Group, and Outside Group. The first region covered the protected territories and included two endorheic lakes: Azhibeksor and Tengiz – both Large (LT) and Small Tengiz (ST) – as well as two small water bodies next to ST. Other regions consisted of 2 to 10 shallow endorheic lakes. Overall, 15 lakes and 29 sampling sites were included in the experiment. The sites were labeled with a lake name or a letter code if the name was unknown. Numbers indicate sites that were located within the same lake. Regions were consistently color coded.

**Figure 1.**
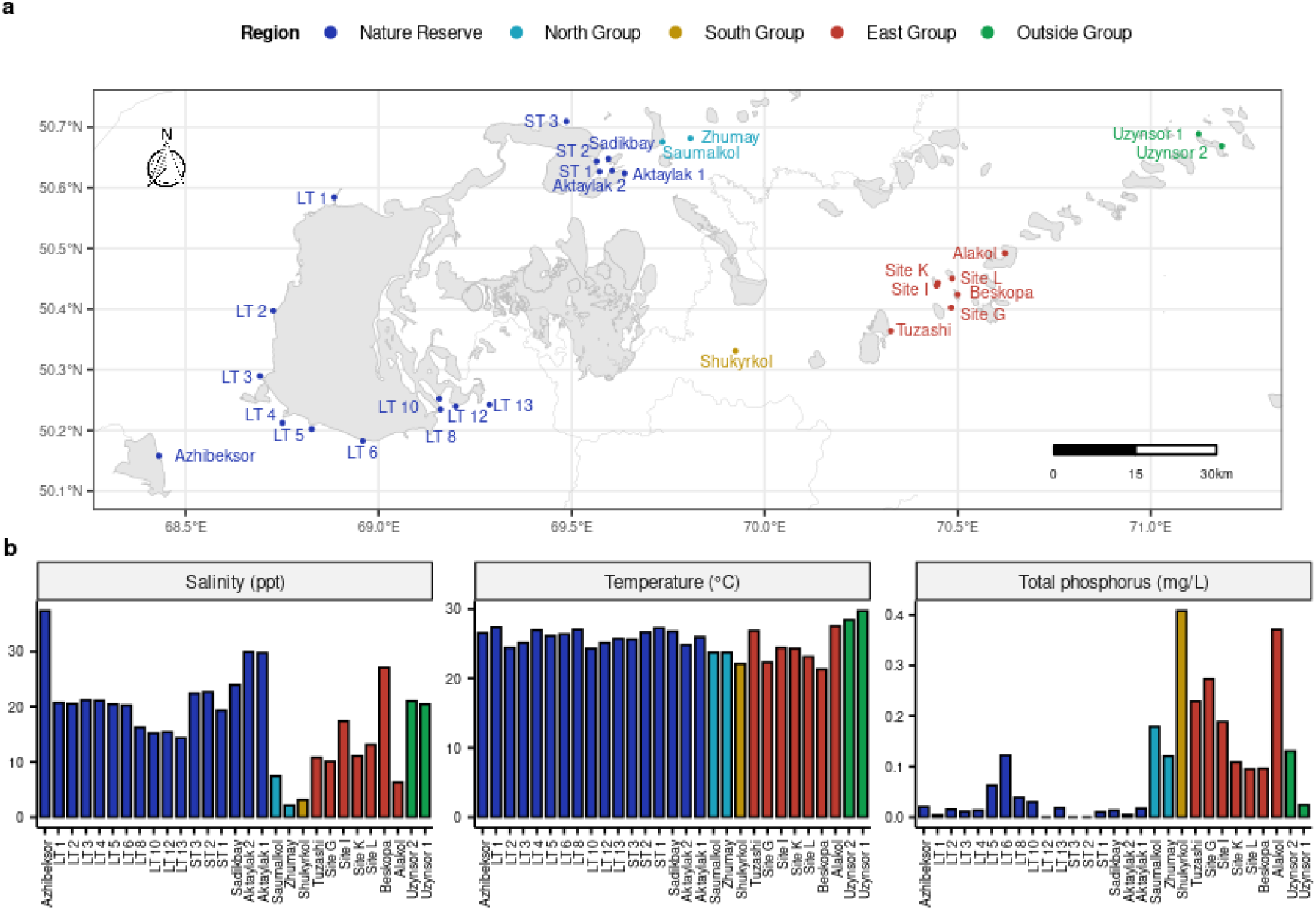
Sampling sites details. (**a**) Geographic location (**b**) and environmental variables: salinity (‰), temperature (=), and total phosphorus (mg/L). Created with use of OpenStreetMap (CC BY-SA 2.0).

### Sample collection and processing

All water samples used for this study were collected in the coastal zone of the lakes during several consecutive expeditions to TKL in July-August 2021. Upon delivery to the laboratory, biomaterial was filtered using a vacuum pump onto the 0.22 µm glass fiber membrane filters (Millipore, USA) and then stored in 50-ml Falcon tubes (BD Biosciences, USA) at −80 °C. The following physico-chemical parameters were recorded for each sample on site: temperature, conductivity, pH, total dissolved solids (TDS), salinity using Cyberscan PC 300 multimeter (Eutech Instruments, Thermo Fisher Scientific Inc., USA) and dissolved oxygen (DO) using a YSI Pro Plus multimeter (Xylem Inc., USA). The total phosphorus content was estimated using protocols by the U.S. Environmental Protection Agency (EPA)^36^.

### DNA extraction, library preparation, and sequencing

DNA was extracted with the PowerWater DNA Isolation Kit (Qiagen, MD, USA) according to the manufacturer’s protocol and stored at −20 °C. The purity and concentration of the DNA were assessed with Nanodrop (Thermo Fisher Scientific Inc., USA).

PCR was performed under standard conditions with Dream Taq Hot Start PCR Master Mix 2X (Thermo Fisher Scientific Inc., USA). The purification step was performed with AMPure XP magnetic beads (Beckman Coulter, CA, USA). The ONT 16S Barcoding Kit SQK-16S024, the Flow Cell Priming Kit EXP-FLP002, and MinION R9 (FLO-MIN106D) were used for library preparation and sequencing (Oxford Nanopore Technologies, Oxford, UK).

Basecalling and demultiplexing were completed using GPU-based Guppy (version 6.4.6, Oxford Nanopore Technologies, UK). Reads were then filtered by length and quality: a range of 1300 -1650 base pairs and a Q-score of at least ten were set as inclusion criteria.

### Taxonomic classification

Taxonomic classification and relative abundance estimation were performed using the Emu algorithm, designed for long and noisy Oxford Nanopore reads^32^. The custom reference taxonomy database was used, which is a combination of rrnDB v5.8^37^ and NCBI 16S RefSeq^38^ downloaded on May 12, 2023. The custom database consists of 19,627 unique species that are represented by 67,931 reference sequences.

### Statistical analysis

Analysis was performed with R version 4.3.0^39^, R Studio version 2023.6.0.421^40^, and the R packages phyloseq v1.44.0^41^ and vegan v2.6-4^42^ were used to handle abundance, environmental, and geographical data. Rarefaction without replacement was performed with the rarefy_even_depth() function from the vegan package. The rarefaction depth was 50,000 reads per sample. Hill diversity indices were chosen as alpha diversity measurements to explore community composition on arithmetic, logarithmic, and reciprocal rarity scales: observed richness (i.e., number of species), Hill-Shannon entropy, and Hill-Simpson concentration index^43,44^. Evenness (J) was calculated with the Pielou’s formula^45^:

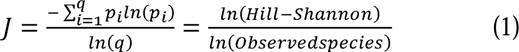

where p_i_ is species relative abundance and q is the number of species.

The correlation between biodiversity and environmental parameters was evaluated based on Pearson’s coefficient. The difference in the composition of the bacterial communities was calculated using Bray-Curtis dissimilarity and then visualized on non-metric multidimensional space (NMDS). The ordination stress value of 0.1 or less was considered satisfactory with a low risk of misinterpretation. In the case of high-stress values, three-dimensional solutions were searched. The final plot was rotated to maximize the variance on the first dimension. Analysis of similarity (ANOSIM) was performed to compare community similarity at different scales (region, lake, site). Mantel test was used to check for correlation between abundance, environmental, and geographical distance matrices^46^. Explanatory power of the environmental and geographical variables on species variation – also called direct gradient analysis – was explored with Canonical Correspondence Analysis (CCA). Partialling out spatial and environmental variation in community structure was performed according to a method described by Borcard and co-authors^47^. The multipatt() function and the group-equalized ‘indicator value’ (IndVal) index from the indicspecies package were used to determine indicator species associated with groups of sites^48^. IndVal is the product of two probabilistic values, called A and B: probability of a site where the species is found to be a member of the site-group and the frequency of the species being found at sites that belong to the site-group, respectively.

## Results

### Geographical and environmental data

Geographical and environmental information of the 29 collection sites (comprising 15 lakes and 5 regions) is shown in **Figure 1** and **Supplementary Table 1**.

### Bacterioplankton community richness and composition

#### Alpha-diversity and community evenness

Based on the species-level classification of 16S sequences, 3290 distinct bacterial species, 1584 genera, 457 families, 180 orders, 83 classes, and 38 phyla were identified across the sampling sites: per-site estimates are given in **Figure 2** and **Supplementary Figure 1.** The taxa were heterogeneously distributed, with the majority of species contributing less than 0.1% to the total bacterial count. The observed richness of lake bacterial communities ranged from 365 (Azhibeksor and Zhumay) to 1026 (ST1) species with a mean of 665 (±173) distinct species per sample, and it was negatively correlated with community evenness (Pearson’s r = −0.39, p-value = 0.042). Hill-Shannon ranged between 38 (ST 2) and 214 (LT 3 and Alakol), with mean of 118 (±46), while Hill-Simpson ranged between 6 (ST 2) and 91 (Alakol) with an average of 40 (±24). The species diversity, expressed in Hill numbers, showed strong linear relationship (R-squared [0.77 - 0.96], p-value < 0.001) with estimates at genus and family levels; goodness of fit dropped significantly (R-squared [0.14 - 0.41], p-value < 0.05) when comparing species and class levels (**Supplementary Figure 2**).

**Figure 2.**
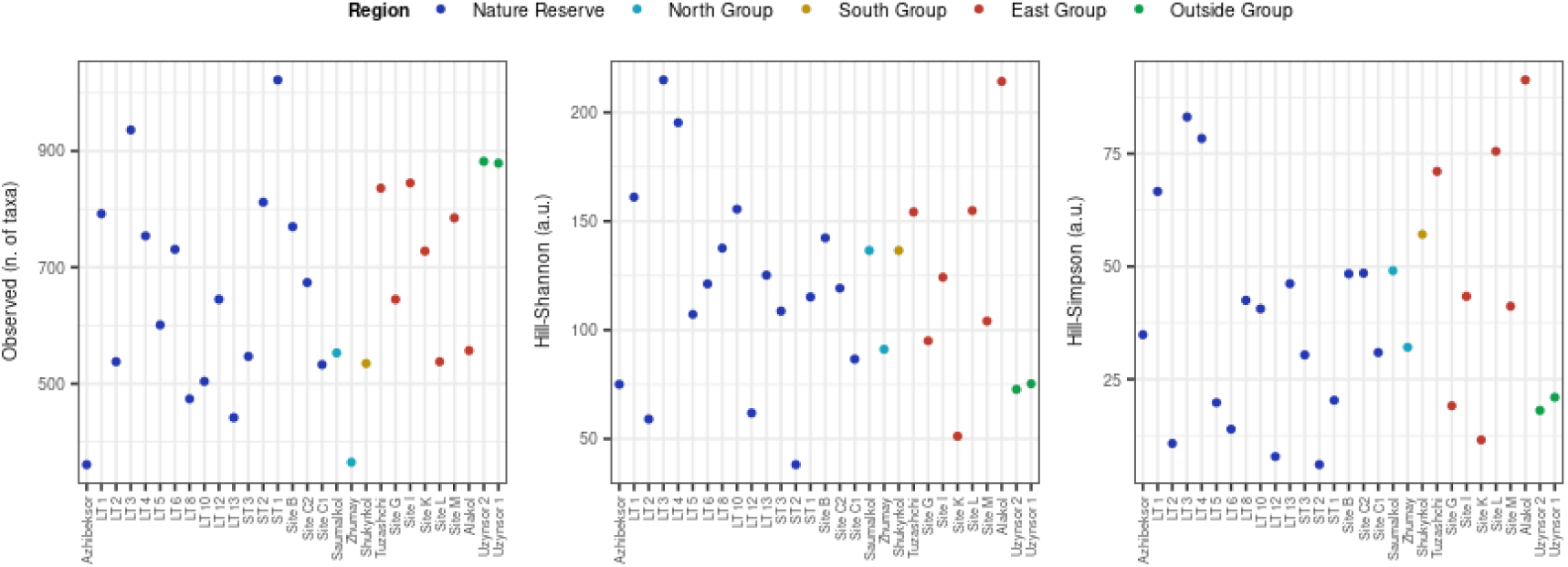
Richness and alpha-diversity estimates for the lake samples: Observed richness, Hill-Shannon, and Hill-Simpson.

Based on Pearson’s correlation test, alpha diversity (observed richness, Hill-Shannon, Hill-Simpson) was not found to be significantly correlated with any environmental variables (salinity, temperature, dissolved oxygen, TP). The small number of sites per region did not meet the minimum requirements for statistical testing, but the visual inspection did not reveal any potential dependence (**Supplementary Figure 3**).

#### Beta-diversity and community composition

The six most abundant bacterial phyla present across all sites were Pseudomonadota, Bacteroidota, Actinomycetota, Cyanobacteriota, Bdellovibrionota and Campylobacterota (**Figure 3**); note that the latter two were previously considered to be a part of the Proteobacteria (Pseudomonadota) phyla.

**Figure 3.**
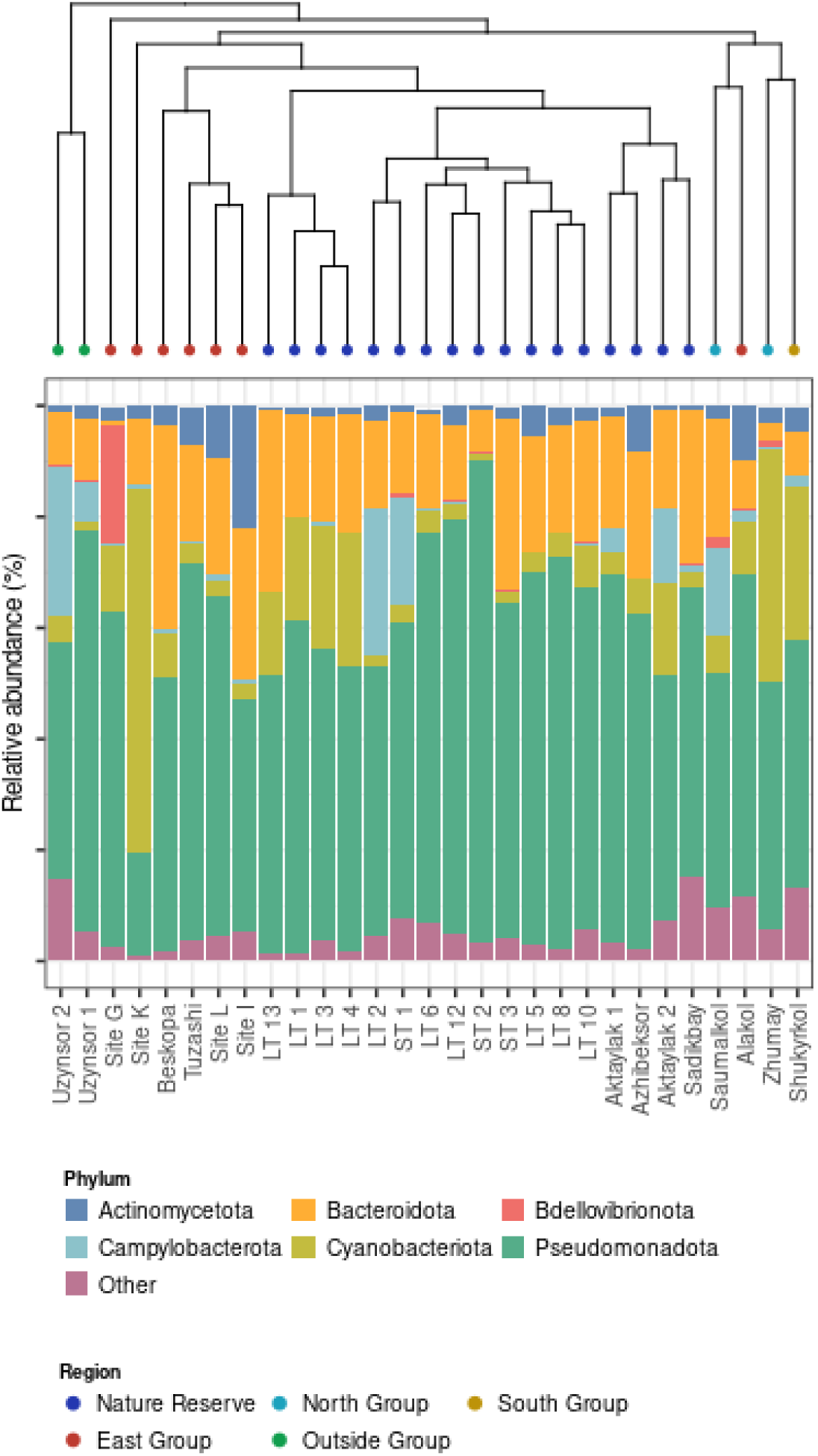
Bray-Curtis-based McQuitty clustering and phylum level composition of the sampling sites. The percentages of the six most abundant phyla are included, the remaining groups are classified as ‘Other’.

The dissimilarity in microbial community composition was well characterized by both clustering and ordination (**Figures 3, 4**). Both techniques identified the Outside Group samples as outliers compared to the other regions. In the Nature Reserve, the Tengiz samples were plotted closely together with the three remaining lakes: Azhiberksor, Sadikbay and Aktaylak. While Azhibeksor was localized somewhat separately, lakes Sadikbay and Aktaylak were associated with the Small Tengiz samples. While Azhibeksor was localized somewhat separately, samples from lakes Sadikbay and Aktaylak were associated with the Small Tengiz samples. The sites from the rest of the regions were more scattered. With a certain degree of regional fidelity, the East Group samples were quite heterogeneous with some resemblance to Nature Reserve (Beskopa), South Group (Alakol), or North Group (Site G). Site K showed high dissimilarity from its group only in clustering output. The Zhumay and Saumalkol sites (North Group), whilst having site-specific bacterial signatures, were more closely related to each other than to all other regions. Notably, sites Zhumay, Saumalkol, Alakol, and Shukyrkol, although coming from different regions and being plotted in a scattered manner (**Figure 4a-b**), were clustered together in a low-salinity (< 10 ‰) cluster (**Figure 3**).

**Figure 4.**
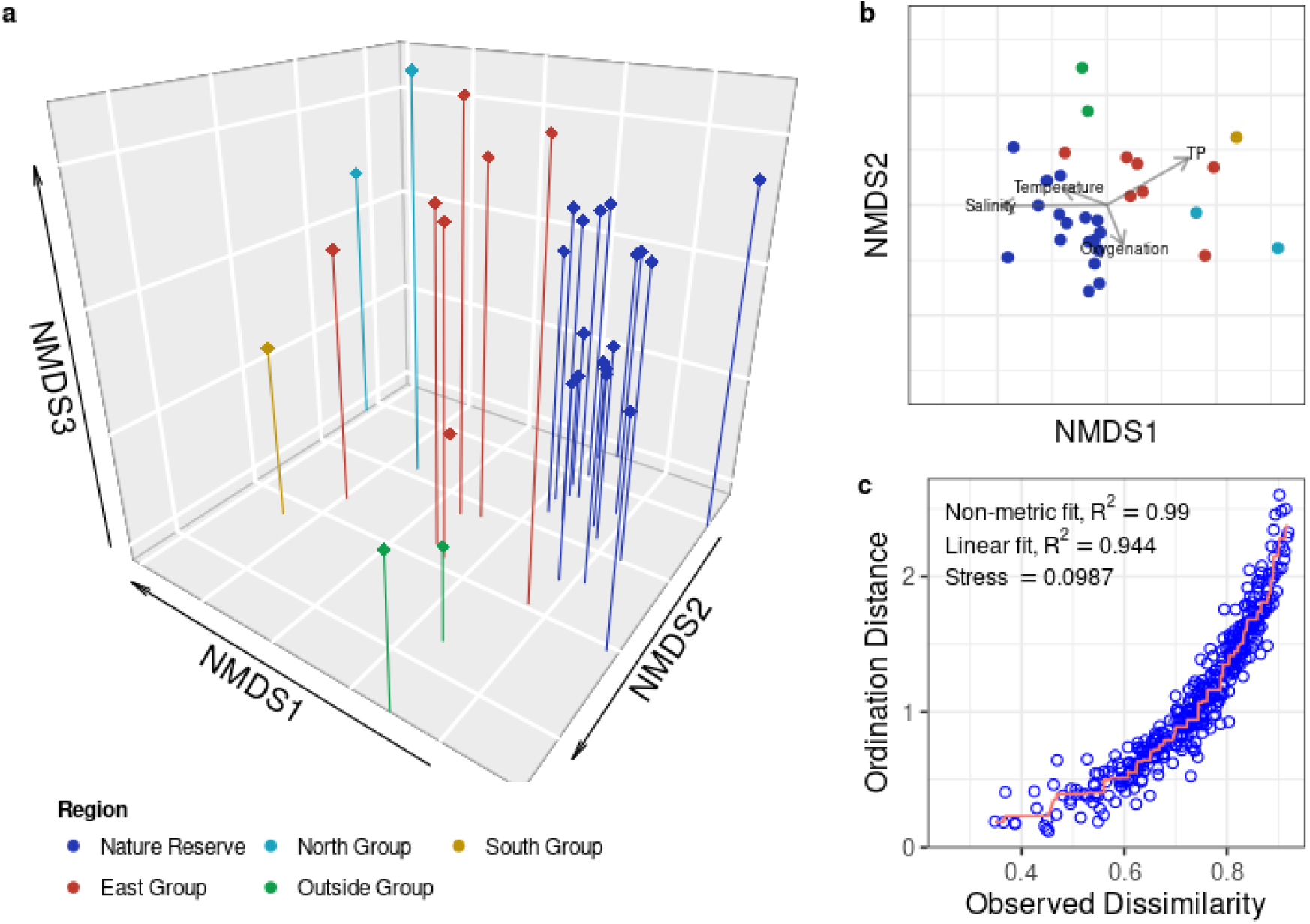
NMDS ordination based on the Bray-Curtis dissimilarity matrix. (**a**) 3D ordination plot, (**b**) 2D representation with fitted environmental parameters, and (**c**) Shepard’s plot.

Focusing on the abundant taxa – defined having relative abundance of > 0.1% at genus level in at least one of the samples – we examined the core microbiome of the lake system. The total taxonomic pool included 597 genera and 1965 species found across the sampling area. Based on the presence-absence data, 127 (21.3%) genera and 138 (7.0%) species constituted the core microbiome in all five regions (**Figure 5**), and even smaller proportions were observed to be present in all 15 lakes (5.7% and 1.7%, respectively, see **Supplementary Table 2**). Almost two-thirds of the lake-wise core species were representatives of Cyanobacteriota – a phylum constituting a relatively modest share of the total community (**Supplementary Table 2**). The core microbiome increased upon exclusion of the low-salinity cluster, with 348 (58.6%) genera and 521 (27.4%) species being shared among the three regions (data not shown). Notably, whilst there was clear regional (and even lakes-wise) heterogeneity in the composition of bacterial species, this dissimilarity was less resolved at the genus level.

**Figure 5.**
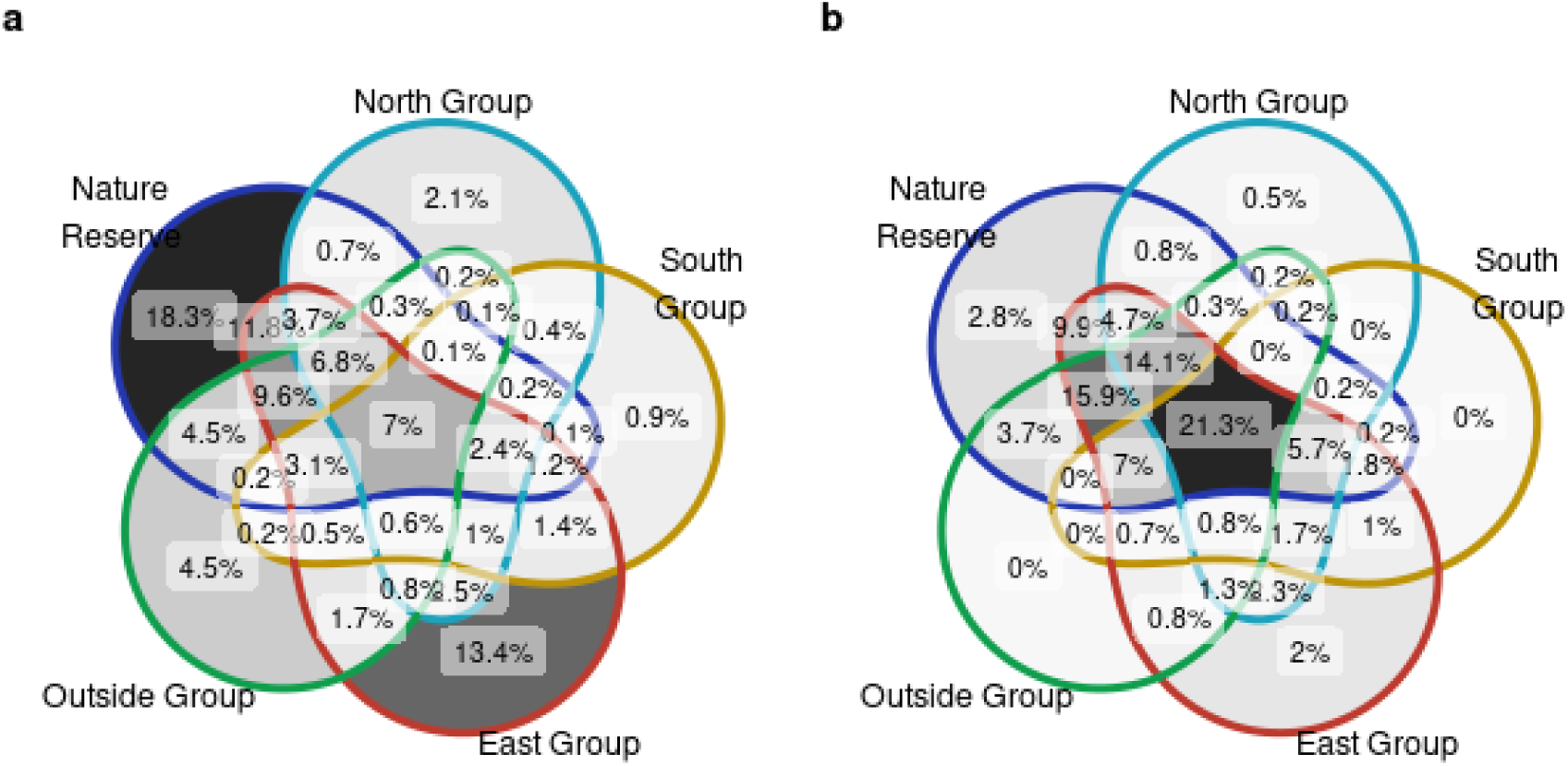
Regional distribution of bacterial taxa (region as a unit of sites): based on (**a**) species (n = 1965) and (**b**) genera (n = 597) presence-absence data.

### Driving factors of microbial diversity and indicator species

#### Geographical patterns in species distribution

Sampling sites located in the same lake region (ANOSIM R 0.8268, p-value < 0.001) or closer to each other (Mantel r = 0.3429, p-value < 0.001) were more alike in terms of microbiome composition. Exclusion of rare taxa did not affect the ability of ANOSIM to resolve the geographical pattern in the remaining community (ANOSIM R 0.8266, p-value < 0.001). To investigate the bacterial taxa that contribute to this pattern, we performed the indicator value analysis. Species showing significant association with region combinations are reported in **Supplementary Table 3**. Among 1965 species, 172 (8.75%) showed significant association to one region, 72 (3.66%) were associated with combinations of two regions, while 67 (3.4%) and 20 (1.02%) were associated with combinations of three and four regions, respectively.

There were 43 species with strong association to the Nature Reserve, the largest yet most homogenous group in terms of microbial composition. Only three species (*Marinomonas communis*, *Roseibacterium beibuensis*, and *Loktanella acticola*) were identified to be both restricted to the region (A = 1.00) and present at all its sites (B = 1.00), and 14 more species exhibited a patchy distribution across the region (0.76 < B < 0.95). Some indicator species were not completely restricted to the region, but appeared in small quantities at other sites (A < 1.00, B = 1.00). Examples of this were the most abundant bacteria *Candidatus Pelagibacter* sp, whose relative abundance ranged between 1.36% and 39.42%, and other less abundant species such as *Kistimonas scapharcae*, *Marinomonas gallaica*, *Marinimicrobium* spp (*M. agarilyticum*, *M. locisalis*), *Neptunomonas phycophila,* and three *Oceanospirillum* spp (*O. beijerinckii, O. multiglobuliferum,* and *O. sanctuarii*). Indicator species accounted for 4.21% to 44.2% of the total bacterial count across the region, with a median of 20.2%.

The second largest region, East Group, represents a cluster of sites with a very heterogeneous community composition: not a single bacterial species was observed in all lakes across the region. First of all, there were three outliers, as suggested by the clustering and ordination results: Site K was dominated by five *Cyanobacteriota* spp (about 25% of the total bacterial community) all of which were a part of the core microbiome; Site G was dominated by *Fluviispira sanaruensis* (about 25% of the reads); Alakol had an overall distinct bacterial profile. Second, many species with moderate fidelity to the East Group were actually associated with a combination of regions, such as East Group & Nature Reserve (e.g., *Pedobacter* spp, *Pseudomonas* spp), East Group & Nature Reserve & North Group (e.g., *Burkholderia* spp, *Marinobacterium ramblicola, Duganella alba, Microbulbifer aggregans*), East Group & Nature Reserve & Outside Group (e.g., *Marivita* spp), etc.

The North Group, although consisting of only two sites, was also quite heterogeneous. When looking at the presence-absence data, we identified 42 species unique to the region; however, there was almost no overlap between two lakes. Thus, 59.5% percent of the taxa were found explicitly in Zhumay, and 35.7% in Saumalkol – most of them had a relative abundance of about 1% or less. Similarly, the indicator value analysis identified only four low-abundance taxa with strong regional association. While Zhumay had a more distinct bacterial profile, Saumalkol had some species in common with neighboring water bodies from the Nature Reserve; *Microbulbifer* spp, for example, were common across the East Group, Nature Reserve, and Saumalkol lake sites.

The Shukyrkol and Uzynsor sites were both the only representatives of their respective regions, hence the inflated number of indicator species (**Supplementary Table 4**), especially those with high regional fidelity (A = 1.00) and frequency (B = 1.00). Yet, considering the fact that there was an average of 15 unique species per sampling site (presence-absence data), this result was expected. Although the Outside Group had many overlaps with other sites, it was mostly set apart due to the low alpha diversity and thus increased abundance of certain species. In fact, the 12 most abundant species accounted for about 50% of the community at Uzynsor sites, of which seven belonged to the region-wise core microbiome, while the remaining five were found in all regions except for the low-salinity cluster.

Across the 31 combinations of site-groups generated by the indicator value analysis, we observed prominent species-level patterns in distribution of bacterial taxa: numerous congeneric indicator species had a strong association with different combinations of lake regions (e.g., *Pedobacter* spp, *Clostridium* spp, *Legionella* spp, *Roseovarius* spp, *Phaeodactylibacter* spp, *Erythrobacter* spp, *Mucilaginibacter* spp, *Flavobacterium* spp) (**Figure 6**).

**Figure 6.**
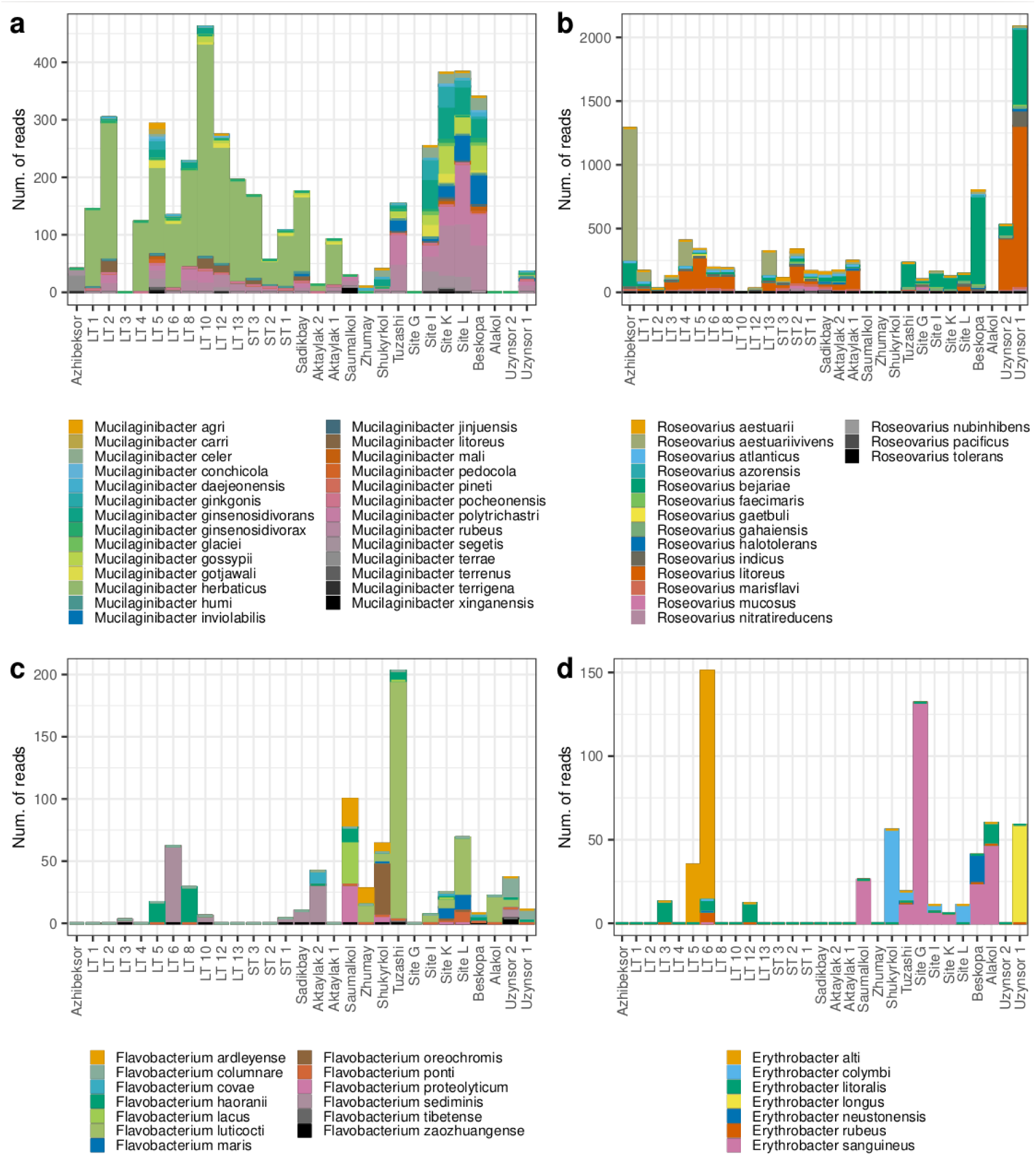
Several congeneric indicator species are showing association with different lake regions. (a) *Mucilaginibacter* spp, (b) *Roseovarius* spp, (c) *Flavobacterium* spp, (d) *Erythrobacter* spp.

#### Partialling out the geographical component of variation

To distinguish between the geographical and environmental factors influencing the bacterioplankton composition in the lakes studied, we focused on the following variables in the CCA model: salinity, total phosphorus, temperature, dissolved oxygen, site region and exact geographical coordinates. Overall, the environmental and geographical parameters explained 50.3% of the total variation (total inertia = 5.12): spatial factor accounted for the majority of the constrained variation with a slight overlap with environmental variables (**Figure 7a**). A large part of the variation remained unexplained.

**Figure 7.**
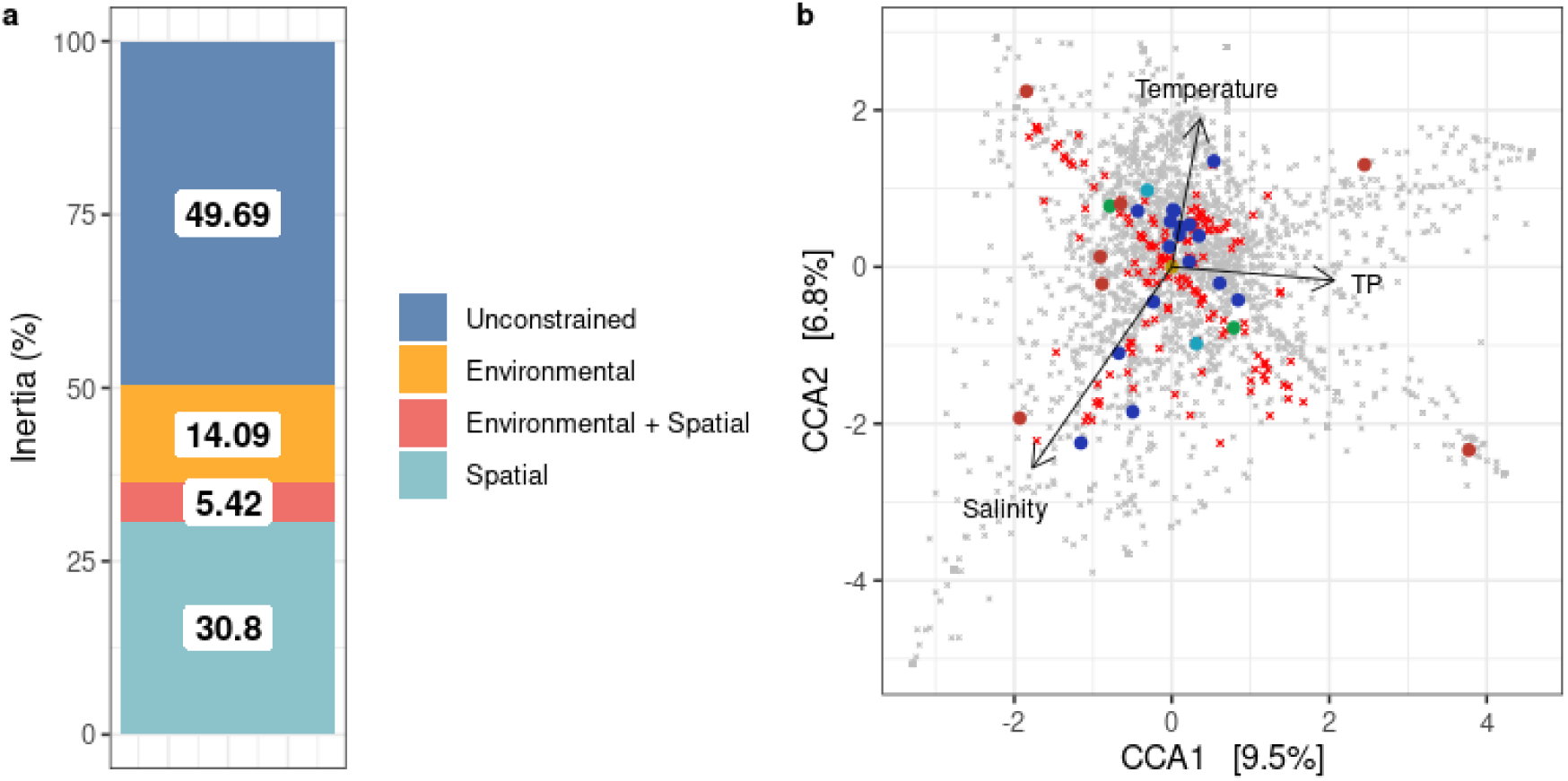
Partialling out components of bacterial species variation. (**a**) Percent of the total inertia explained by environmental parameters and spatial structure. (**b**) Partial CCA triplot of the Bray-Curtis matrix, constrained by the environmental matrix, with removed effect of geographical matrix; region-specific species are shown in red (**Supplementary Table 3**), the remaining species in gray.

Removal of the geographical effect practically eliminated the differences between Outside Group, South Group, and the Tengiz sites, but highlighted distinct communities of lakes adjacent to Tengiz (Azhibeksor, Sadikbay and Aktaylak) and the heterogeneity of East Group lakes (**Figure 7b**). Even though the effect of spatial association was constrained on the graph (**Figure 7b**), some indicator species with strong regional preference (red) show distribution along the environmental gradient, i.e., salinity.

#### Distribution of bacterial species along environmental gradients

Mantel tests indicated a significant correlation between environmental parameters and microbial community composition. Three major factors affecting community dissimilarity were salinity (Mantel r = 0.52, p-value < 0.001), TP (Mantel r = 0.48, p-value < 0.001), and water temperature (Mantel r = 0.40, p-value < 0.001): sites with similar salinity, TP, and temperature tended to have more similar microbial composition. Dissolved oxygen, on the other hand, did not significantly correlate with bacterial abundances (Mantel r = 0.03, p-value = 0.38).

We implemented the same method as in the section above (IndVal) to identify species specific to the low-salinity cluster (Zhumay, Saumalkol, Shukyrkol, and Alakol), which was previously highlighted by the clustering method. Notably, the low-salinity cluster covered two geographic regions (North and South Groups), implying that region- and lake-specific indicator species are as likely to be determined by salinity as by geographical factor. The list of 22 species associated with at least two of the sites (A = 1.00, B ≥0.50) is displayed in **Supplementary Table 4**. The most prominent examples were the congeneric species with similar response to environmental selection, such as *Limnohabitans* spp (*L. planktonicus, L. parvus,* and *L. radicicola*) and *Polynucleobacter* spp (*P. asymbioticus, P. difficilis,* and *P. cosmopolitanus*). In some cases, however, individual species responded differently to environmental conditions: *Algoriphagus* spp, *Rhodoluna* spp, *Pseudohongiella* spp (**Figure 8**).

**Figure 8.**
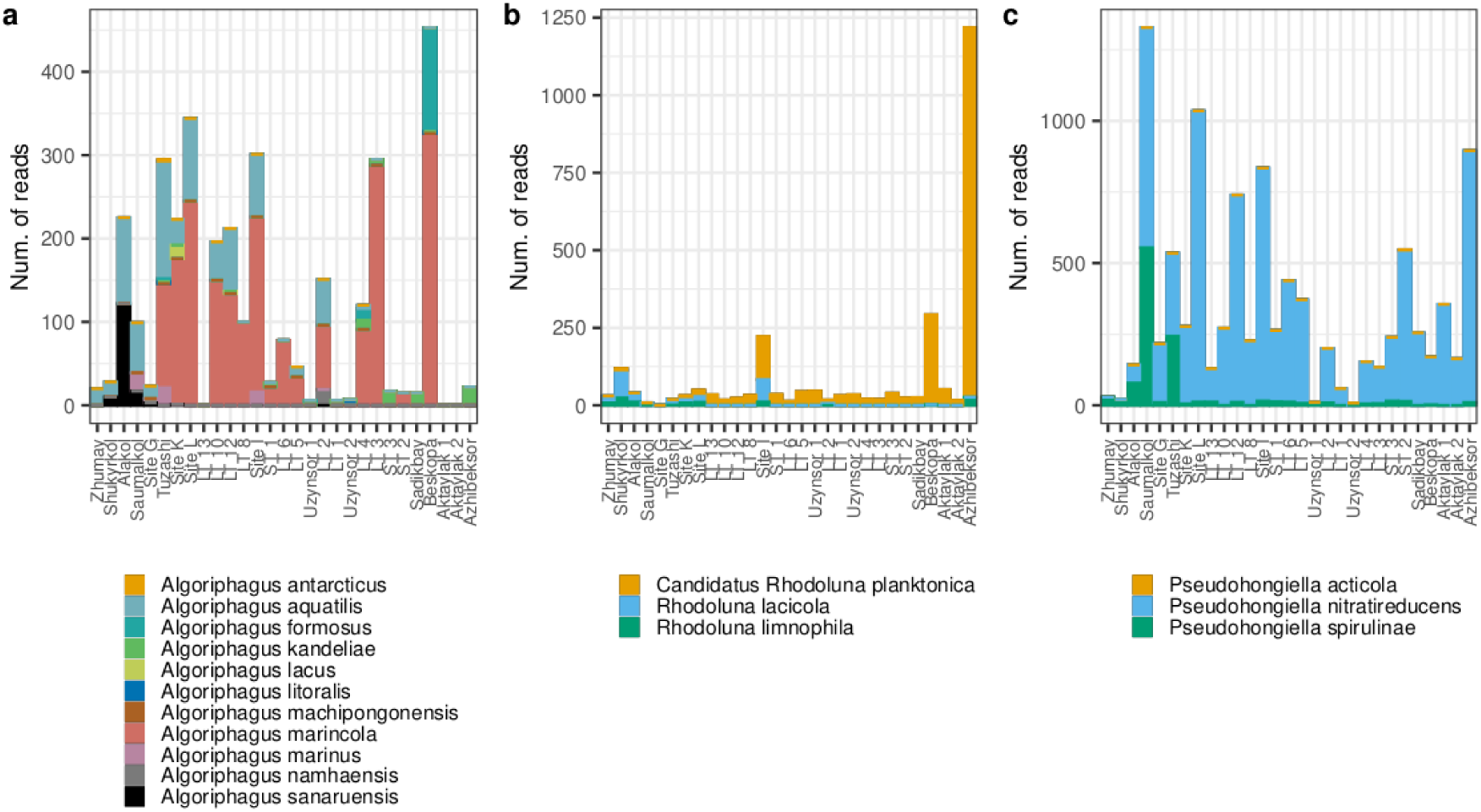
Differential abundance of congeneric indicator species in response to salinity: (a) *Algoriphagus* spp, (b) *Rhodoluna* spp, (c) *Pseudohongiella* spp. Sites are displayed in the order of increasing salinity.

Apart from qualitative differences, the salinity gradient also exerted a quantitative effect on the community profile. We evaluated the relationship between the relative abundance of bacterial species and salinity percentage with the Pearson’s product-moment correlation. Out of the 117 indicator species associated with the Nature Reserve (or its combination with other regions), 52 bacterial species correlated significantly (Pearson Product-Moment Correlation, p-value < 0.05) with salinity. 24 species, mainly represented by the genera *Marinimicrobium*, *Marinobacterium*, *Marinomonas*, *Neptunomonas*, *Oceanospirillum,* and *Pseudomonas* (*Gammaproteobacteia*), were positively correlated with salinity (**Supplementary Figure 4**). The remaining species, members of *Burkholderiaceae, Chitinophagaceae, Oxalobacteraceae, and Sphingobacteriaceae* families, correlated negatively with salinity (**Supplementary Figure 5**). Among the taxa associated with the East Group, 16 species were found to correlate positively with salinity (**Supplementary Figure 6**); many of the taxa overlapped with those from Nature Reserve, e.g., *Marinobacterium* spp, *Neptunomonas* spp, *Pseudomonas* spp, showing consistent trend along the salinity gradient but different levels of relative abundance (**Figure 9**).

**Figure 9.**
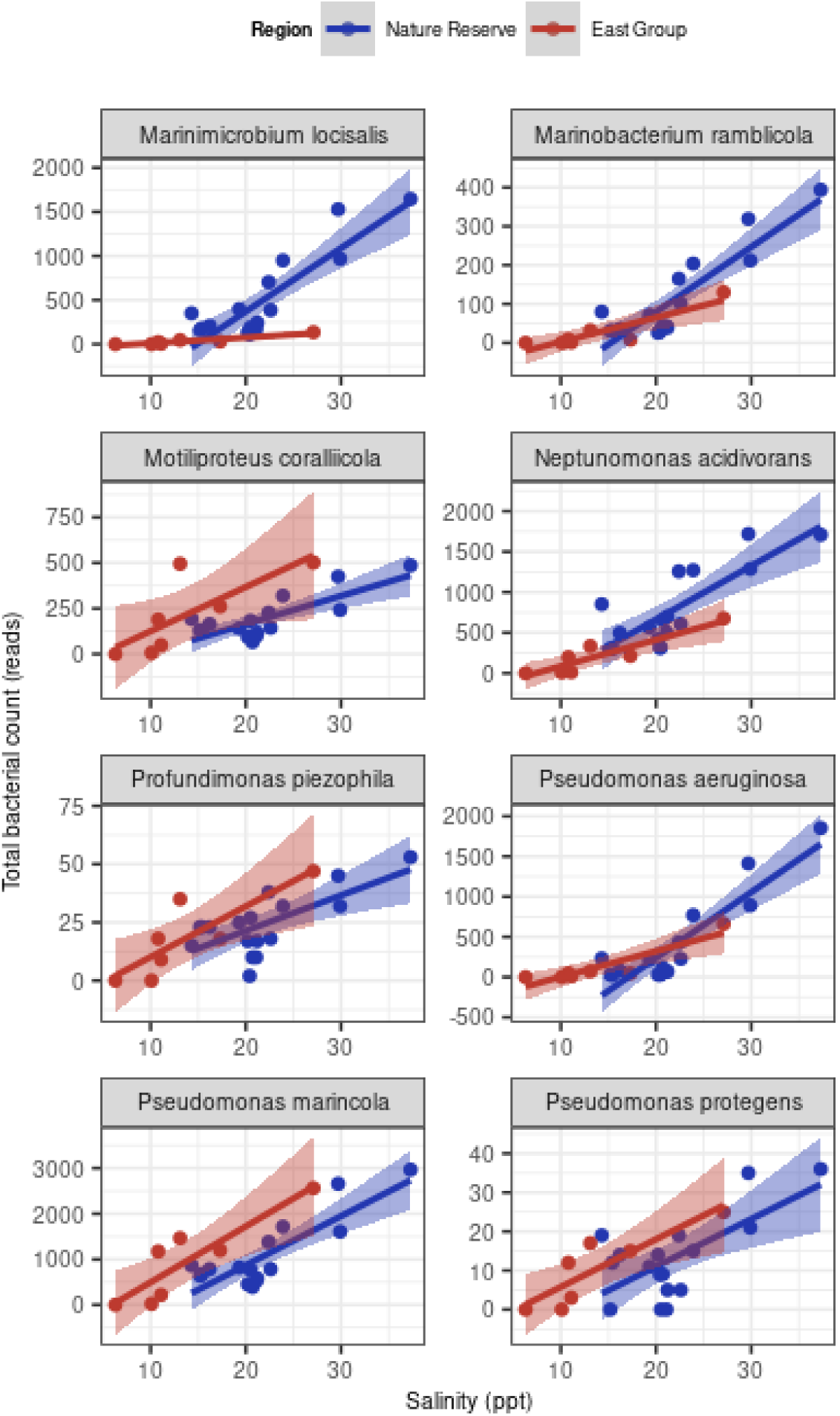
Relationship between salinity and the relative abundance of indicator species common for the Nature Reserve and East Group.

## Discussion

### Alpha-diversity is not a sufficient community descriptor

The average number of unique species per sample (S) scaled with the number of reads (N = 50,000) at a rate of ∼ 0.6 (i.e., S ∼ N^0.6^), which was slightly greater but close to the expected range of [0.25 - 0.5]^49^. The negative correlation between the number of species and evenness complied with the diversity scaling law, i.e., samples with a high number of observed species were actually inhabited by a small number of abundant bacteria and many rare taxa^49^. Hill numbers of higher order lend more weight to the relatively abundant taxa. Hence, a decrease in “the effective number of species” was observed. In some cases this decrease was drastic, e.g. LT2, LT12, ST2, and Site K (**Figure 2**), suggesting that most of the taxa determined at a given site were rare. In fact, 25-40% of the reads from these samples were represented by a single taxon; yet, long-read HTS allows to recover three to five hundred rare taxa per site. However, while effectively resolving bacterial diversity, species-level classification does not provide significant advantage over genus-level studies at this stage. Overall, alpha-diversity estimators, albeit they provide a fair overview, were not useful for disentangling the biogeographical patterns of bacterioplankton communities since the relationship with environmental variables or geography was not evident.

### Shallow endorheic lakes show unusually low bacterioplankton dispersal rates

Our results demonstrate that while covering large spatial and environmental scales, the microbial community at the Tengiz sites is relatively homogeneous. The inter-lake variability has much higher magnitude. Many studies have previously highlighted that bacterial dispersal rates are affected but not significantly limited by geographical scales, and that it is common for water bodies located several thousand kilometers from each other to share a large portion of their microbiome^50–52^. We, however, observed that an unusually high proportion of variation could be explained by the geographical distance between sites and their location (region) on a scale <200 km (**Figure 7**), and the percentage of microbial taxa shared was only 7% across all five regions and 27.4% across saline lake regions, compared to >85% found by Van der Gucht and coworkers (2007). There might be three facets to this observation of geographical importance.

The first facet is skeptical and claims that the explanatory power of geographical factors is attributed to a variable with regional differences that we did not take into consideration in our analysis. Anthropogenic factors, such as proximity to farmlands or villages, could potentially explain the relative homogeneity of the Nature Reserve region (restricted access area) compared to the rest of the regions studied. Regional preferences of phyto- and zooplankton, fish, migratory and nesting birds populations^34,53^ might be reflected in the microbial composition as a result of biotic interactions. Lastly, additional spatially autocorrelated abiotic interactions not considered in the present study could play a role.

The second explanation is that high heterogeneity of lakes bacterial communities is a specific characteristic of the studied ecosystem. In arid climates, shallow endorheic lakes are shaped by the flooding and desiccation dynamics, and exhibit frequent changes in temperature and salinity, sometimes turning into ephemeral water bodies. Such unstable inland lakes systems have been previously reported to exhibit high genetic diversity and heterogeneity^54^.

The third explanation may relate to in-lake variability and can be based on the heterogeneous physiological characteristics of different bacterial species and persistence of bacterial assemblages across spatial scales. Shallow lakes usually lack stratification and appear in two different ecological states depending on submerged macrophytes^55^. In contrast to smaller habitats, large lakes such as Lake Tengiz (e.g., lakes Taihu^56^ and Dongting^57^, China) exhibit significant environmental gradients and may harbor both ecological states within the same lake. It puts Lake Tengiz apart from small lakes that were sampled one site in each lake.

The fourth facet is methodological and emphasizes the role of higher taxonomic resolution. In a meta-analysis study, Hanson and colleagues (2012) have concluded that even though spatial structure has been rarely highlighted as a major community driver in previous microbiome studies, a positive trend has been observed between the increasing precision of taxonomic classification and a relative effect of the spatial component. According to our observations, the species-level classification achieved with the long-read sequencing indeed allowed us to identify dispersal patterns not resolved previously when classification was limited by higher taxonomic levels, such as genera and families.

### Salinity is the major environmental gradient driving microbiome composition

Even though salinity did not correlate significantly with alpha diversity estimators, we identified it to be the main environmental variable driving microbial composition. This is in line with the global patterns of microbial distribution^20^ as well as with results of studies focused on saline lakes and estuaries^58–60^.

The most drastic shift in microbiome composition occurred above the salinity threshold of approximately 10‰, which contrasted lakes Zhumay, Alakol, Shukyrkol, and Saumalkol (low-salinity cluster) with other sites, this being even more striking as these four sites are located in different regions of the TKL. The two highly abundant *Betaproteobacteria* shown to be either restricted to or prevalent in low-salinity lakes (< 10‰) were the genera of free-living freshwater bacteria *Polynucleobacter* and *Limnohabitans*^61^. In addition, several indicator species from *Alphaproteobacteria* (*Rhodobacter* spp, *Caulobacter* spp, and *Tabrizicola* spp), *Actinomycetes* (*Rhodoluna lacicola*), *Bacteroidota* (*Aquirufa* spp, *Algoriphagus sanaruensis*), and *Cyanobacteriota* (*Planktothrix agardhii*) are also reported for freshwater habitats^62–66^. Besides these planktonic freshwater taxa, the indicator value analysis demonstrated presence of a high number of shared soil-derived bacterial groups. On the one hand, inclusion of soil bacteria via dust or sediment cannot be avoided when taking coastal samples; however, it might also indicate temporal desiccation of lakes; for example, as reported by the Association for the Conservation of Biodiversity of Kazakhstan, Zhumay (one of the lakes studied in this work) was completely dried out between years 2010 and 2013, until its restoration via snow retention^67^. Such shallow ephemeral lakes are likely to have representatives (potentially dormant) of biocrust communities and exhibit overall high heterogeneity in diversity estimations^68^.

Even though it is common for closely related taxa to exhibit similar ecological preferences, implementation of the long-read sequencing and species-level metagenomics enables resolution of divergent biogeographical patterns even for congeneric species. For example, distribution of *Algoriphagus* spp across sampling sites closely followed the optimum salinity conditions described in the literature: *A. sanaruensis* was associated with the low-salinity cluster; *A. aquatilis* was transitional for the low-salinity and East Group sites; *A. marincola* was distributed across sites with salinity above 10‰, and *A. kandeliae* had a preference for high salinity sites (> 20‰)^69–72^.

As described in the current study, the bacterial profile for sites with salinity > 10‰ was less uniform; despite the overlap in salinity ranges between Nature Reserve, East Group, and Outside Group, regions only shared a handful of species. In the first region, large portion of the microbiome was represented by *Gammaproteobacteria*, mainly from the *Marinimicrobium*, *Marinobacterium*, *Marinomonas*, *Neptunomonas*, *Oceanospirillum,* and *Pseudomonas* genera, all of which are halotolerant and halophilic bacteria, naturally showing a positive correlation with salinity^73,74^. Majority of these species were also found in the East Group sites but in much smaller quantities than would be predicted based on salinity, which could imply a potential source of limitation to their dispersal. The only exception with wider dispersal was *Pseudomonas* spp, which were homogeneously spread across both regions with respect to salinity.

## Conclusion

This is the first study that provides a detailed, species-level characterization of environmental microbiomes with insights into the biogeographical patterns in the bacterial diversity of 15 shallow endorheic lakes. We highlight the potential advantages of the implementation of nanopore-based long-read sequencing for high taxonomic resolution of bacterial diversity. Our findings indicate that the Tengiz-Korgalzhyn Lakes system is extremely diverse, featuring more than 3,000 bacterial species. The microbial communities in the area are greatly influenced by biogeographical patterns such as selection and dispersal processes. Environmental selection in the sampled lakes was mostly governed by salinity, serving as both ecological threshold and an environmental gradient. The dispersal processes are greatly limited by connectivity of the lakes and their position in the landscape, resulting in high heterogeneity among the different lakes and regions. Species-level classification is important in establishing ecological as well as spatial structures in bacterioplankton composition and abundance. The detailed mapping of the lakes’ microbiome provides a foundation for further genomic and functional investigations of the major bacterial players in the rapidly changing aquatic ecosystems.

## Acknowledgments

We are thankful to members of the Tengiz-Korgalzhyn Expedition 2021 (Veronica Dashkova, Aidyn Abilkas, Aleksander Koshkin, Kanat Samarkhanov, and others) for their contribution to the sample collection and initial hydrochemical characterization. We acknowledge the assistance and support of Kanat Baigarin in organizing the trip, Massimo Pindo for his insights regarding lab protocol design, and Anne Mette Paulsen for help with English editing. We acknowledge the support of NPO Young Researchers Alliance and Nazarbayev University Corporate Fund “Social Development Fund” for the grant under their Fostering Research and Innovation Potential Program.

## Funding

This research was funded by Ministry of Science and Higher Education of the Republic of Kazakhstan, grant number AP14872028 to N.S.B., and grant number AP14869915 to I.A.V., by the TÜBITAK program BIDEB2232 (project 118C250) and the Carlsberg foundation to E.J.

## Authors’ contributions

P.L. performed the research, analyzed the data, wrote an original draft, reviewed and edited the manuscript. A.M. performed the research, reviewed and edited the manuscript. G.N. contributed in analysis of water samples, reviewed and edited the manuscript. A.C. contributed to study design and data analysis, reviewed and edited the manuscript. E.J. and I.A.V. contributed to the funding, experiment design, supervision, revision, and edition of the manuscript. C.D. and N.S.B. supervised the whole project, wrote and modified the manuscript. All of the authors reviewed the manuscript.

## Competing interests

The authors declare no competing financial interests.

## Data Availability Statement

The datasets generated during and/or analyzed during the current study are available in the NCBI’s SRA repository with BioProject ID PRJNA1045017.

## Figure Legends

**Supplementary Figure 1.** Alpha-diversity and evenness of lake samples at different taxonomic levels; (A) Observed richness, (B) Hill-Shannon, (C) Hill-Simpson, (D) Pielou’s evenness index.

**Supplementary Figure 2.** Relationship between diversity estimates at different taxonomic levels: **(A)** genus versus species; **(B)** family versus species; **(C)** class versus species. R is Pearson’s coefficient.

**Supplementary Figure 3.** Hill’s diversity indices (Hill-Shannon and Hill-Simpson) of sites across regions.

**Supplementary Figure 4.** Species associated with the Nature Reserve that show positive correlation with salinity.

**Supplementary Figure 5.** Species associated with the Nature Reserve that show negative correlation with salinity.

**Supplementary Figure 6.** Species associated with the East Group that show positive correlation with salinity.

**Supplementary Table 1.** Geographical and environmental lake details.

**Supplementary Table 2.** Region-wise core microbiome: species presence-absence data.

**Supplementary Table 3.** Region specific bacterial species sorted by test statistic

**Supplementary Table 4.** Bacterial species restricted to the low-salinity cluster lakes

## Notes

### Competing Interest Statement

The authors have declared no competing interest.

